# Resolving Heterogeneous Mechanical Domains via Physics-Aware Deep Clustering of Single-Molecule Force Spectroscopy Data

**DOI:** 10.64898/2026.08.31.748330

**Authors:** Cailong Hua, Yiyuan Zhang, Vinitendra Singh, Rebecca A. Walsh, Joseph Vavra, Joseph M. Muretta, James M. Ervasti, Murti V. Salapaka

## Abstract

Many biological processes rely on mechanical forces, with protein molecules acting as key mediators. Understanding how proteins respond to mechanical stress is essential for conditions including cardiomyopathy and muscular dystrophy. Natural proteins such as dystrophin and utrophin are composed of heterogeneous folding domains with distinct mechanical properties; deciphering domain-level behavior provides insights into disease mechanisms and informs therapeutic strategies. Single-molecule force spectroscopy (SMFS) enables probing the mechanical properties of entire proteins, yet current approaches struggle to identify heterogeneous folding domains, particularly without prior knowledge. Here, we present the first automated framework to identify heterogeneous folding domains in SMFS data, applying both existing clustering methods and a novel physics-aware deep clustering architecture, LatentUnfold. LatentUnfold learns complementary latent representations from force magnitude and the force-extension physical relationship through dual autoencoders, jointly optimized for clustering assignments. We apply our framework to experimental SMFS data collected from a synthetic two-domain protein (ddFLN4–Titin I27) as well as natural protein constructs of dystrophin and utrophin, with Monte Carlo simulated datasets serving as controlled validation. For the synthetic protein, we recover mechanical properties consistent with previously reported values for each domain. For the natural proteins, we uncover two mechanically distinct domain populations—corresponding to the N-terminal domain and spectrin-like repeats—with differences in both unfolding force and contour length increase, and reveal different unfolding order between them for the first time. This work enables domain-level biological inference, overcoming prior limitations that relied on averaging and overlooked heterogeneity, thus advancing the understanding of mechanical behavior in protein unfolding.

## Introduction

Mechanical forces play a critical role in many biological processes, such as transcription, translation, cell locomotion, protein folding and unfolding, and intracellular information transfer. ^1,2^ Protein molecules, the functional building blocks of life, play essential roles in these processes via regulating mechanical forces. Understanding how protein molecules respond to mechanical forces offers valuable insight into their functional roles and development of treatments for a range of debilitating and fatal conditions. ^3–6^ For example, mutations in the titin gene can compromise its mechanical tension response, leading to dilated cardiomyopathy; ^7,8^ characterizing mechanical properties of shock-absorbing muscle proteins, dystrophin and utrophin, will better inform on their function in vivo, which could lead to an effective therapy for Duchenne Muscular Dystrophy, a fatal disease affecting 1 in 5000 births. ^9,10^

Protein molecules are composed of folding domains which are compact globular structures with hydrophobic cores and hydrophilic surfaces. ^11,12^ Importantly, domains in natural proteins are often heterogeneous, exhibiting distinct functions and mechanical properties. ^13,14^ For example, in dystrophin (Figure 5), the N-terminal domain anchors the protein to actin filaments beneath the sarcolemma, while the spectrin-like triple-helix repeats are hypothesized to unfold in response to mechanical stress. Therefore, investigating the mechanical properties of individual domains under forces can enhance our understanding of disease mechanisms and support the development of therapies. ^15^

Single-molecule force spectroscopy (SMFS) is a well-established method that directly probes structural changes of macromolecules under mechanical force. Advances in instrumentation—such as atomic force microscopy (AFM) ^16^ and optical tweezers ^17^—have enabled SMFS measurements with force sensitivity in the femtonewton to piconewton range and nanometer spatial resolution. However, interpreting SMFS data is challenging, particularly when dealing with proteins with multiple, heterogeneous domains. Natural protein molecules with heterogeneous domains are typical, but current methods are limited to mechanical molecular fingerprints, which are well-characterized protein molecules with known structural or mechanical properties, ^18^ to confirm the presence of single molecules and identify specific domain unfolding events. However, these methods can not be applied to natural proteins comprised of multiple heterogeneous domains. ^19,20^ Moreover, incorporated molecular fingerprints can influence the native behavior of the studied protein, which may result in misleading experimental outcomes. ^21^ In the absence of fingerprints, polymer elasticity models, such as the worm-like chain (WLC) model, ^22^ are employed to fit individual unfolding events, ^23^ but this approach requires domain expert knowledge and does not scale to the thousands of force curves typically collected in an SMFS experiment.^6^ These limitations, coupled with the therapeutic potential of such insights, motivate the development of automated, data-driven tools for identifying heterogeneous domains in natural proteins.

We first adapted a range of existing clustering approaches to the problem of domain identification in force-extension curves, including K-Means and K-medoids with dynamic time warping (DTW), ^24^ autoencoder-based methods using MLP and CNN architectures, ^25^ and deep clustering frameworks such as deep embedded clustering (DEC) ^26^ and deep temporal clustering representation (DTCR). ^27^ While these methods have proved effective for discovering structural patterns in unlabeled time series data, ^28^ clustering unfolding segments from force-extension curves is inherently challenging due to several factors: (1) SMFS experiments exhibit low success rates of single-molecule capture, necessitating the collection of thousands of force curves (typically 2000–5000) to obtain statistically meaningful inferences; ^6^ (2) force curves often contain noise, nonspecific adhesion, and detachment artifacts, complicating the identification of true domain unfolding segments; and (3) often proteins under study possess unknown or complex domain architectures with no prior characterization, making manual annotation by domain experts ^12^ infeasible and error-prone. It is difficult to do manual visualization, which needs expert knowledge, introduces subjectivity, and lacks scalability.

To overcome these challenges, we further developed LatentUnfold, a physics-aware deep clustering framework specifically designed for SMFS data. LatentUnfold employs two complementary autoencoders: a force autoencoder that captures features from force magnitudes, and a relationship autoencoder that models the force-extension relationship informed by polymer elasticity models such as the WLC model. ^22^ These representations are jointly optimized for clustering through a gated fusion mechanism and spectral K-means loss. We conducted experiments on four protein molecules (Figure 5): the synthetic two-domain protein ddFLN4–Titin I27, composed of two well-characterized domains—ddFLN4, the fourth immunoglobulin-like domain from Dictyostelium discoideum filamin, and Titin I27, the 27th Ig-like domain of human titin^1^— and three natural protein constructs spanning the N-terminus (NT) through spectrin-like repeat (SLR) 3: DysN-R3 from dystrophin, and insect UtrN-R3 and bact UtrN-R3 from utrophin, where the two utrophin constructs differ in expression system. ^6,29^ For all three natural constructs, the NT domain is presumed to be structurally distinct from the SLRs, containing roughly twice the number of amino acids. ^10^ Monte Carlo simulated data with known ground truth labels served as controlled validation. For ddFLN4–Titin I27, we successfully separated the two domain types and recovered mechanical properties consistent with previously reported values. For the natural proteins, we uncovered two mechanically distinct domain populations with different unfolding forces and contour length increases, corresponding to the NT domain and spectrin-like repeats, and revealed unfolding order between them for the first time.

This work represents the first reported effort of automating the identification of heterogeneous domains in SMFS data. It provides insight into the mechanical behavior of protein unfolding and highlights the value of domain-level analysis. Our methods would transform the understanding of protein mechanics by revealing how specific structural segments contribute to the unfolding pathway, rather than just the protein as a whole. Previously, a lack of such a facility has limited biological inferences to be drawn in an average sense without accounting for the impact of heterogeneity which is a substantial limitation.

## Results

### Constant Speed Experiments of Heterogeneous Unfolding Domains

In AFM-based SMFS experiments a protein molecule of study is tethered between the tip of micro-cantilever and the substrate. When the cantilever is retracted from the substrate at a constant speed, the force *F* experienced by the cantilever is calculated by *F* = *kd*, where *k* is the spring constant of the cantilever and *d* is the deflection of the cantilever tip. The force *F* acts as a proxy for the tensile force applied to the protein molecule, while the extension *X* is the protein end-to-end distance (Figure 1a), resulting in force-extension curve. The example shown in Figure 1e corresponds to a single protein molecule with two different types of domains, labeled as type 0 and 1. As the cantilever retracts, the first force drop reflects the rupture of nonspecific adhesion between the tip and the substrate. ^30^ Subsequent three force drops are caused by the sequential unfolding of three folded protein domains under mechanical tension (Figure 1b–d). Finally, the protein detaches either from the cantilever tip or the substrate, resulting in the final force drop. ^3^

**Figure 1:**
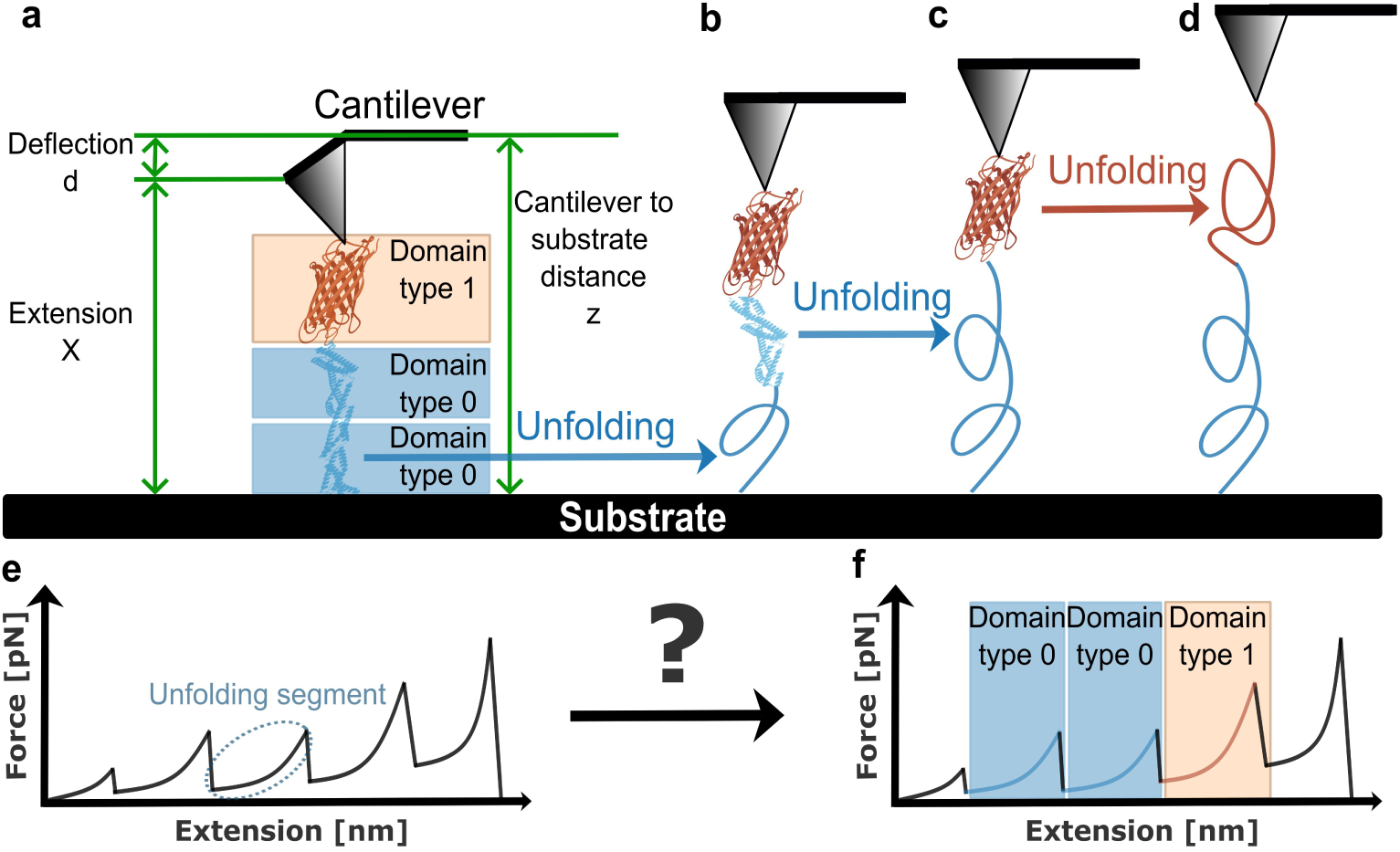
Illustration of AFM-based SMFS. (a) Schematic of the experimental configuration with the protein molecule composed of three folded domains. The deflection *d* and the cantilever–substrate distance *z* are measured during pulling. (b–d) Illustration of sequential unfolding of three domains, including two type 0 domains (blue) and one type 1 domain (red). (e) Representative force-extension curve with the unfolding segment indicated by a dashed oval. (f) Matching unfolding curve segments to domain types 0 and 1, enabling identification of heterogeneous domains.

The force-extension curve (Figure 1e) reflects the unfolding process of proteins composed of multiple heterogeneous domains. Every domain unfolds with its own characteristic force-extension response, termed as an unfolding segment. The transition from Figure 1d to Figure 1e illustrates the goal: resolving the curve into domain-specific unfolding segments—type 0 and type 1. It is crucial to identify which unfolding segment corresponds to which domain type and further deepen the understanding of the biological function of the protein. Therefore, there is a pressing need for robust, automated methods capable of identifying heterogeneous domains from noisy, unlabeled force-extension curves.

### Automated Clustering of Heterogeneous Unfolding Domains

We applied both the existing clustering methods described in the Introduction—K-Means and K-medoids with DTW, autoencoder-based approaches, and deep clustering frameworks (DEC and DTCR)—as well as our novel physics-aware architecture, LatentUnfold, to identify heterogeneous domains from SMFS data. As illustrated in Figure 2, LatentUnfold employs two complementary autoencoders to capture distinct aspects of unfolding segments. The force autoencoder is trained exclusively on force magnitudes; extension values are omitted because identical domains can unfold at different points along the trace, yielding disparate extension readings that would confound representation learning. However, force alone cannot fully describe the mechanical behavior of protein molecules, which is governed by the physical relationship between force and extension. The relationship autoencoder therefore learns from normalized force–extension pairs, embedding the underlying force-extension relationship governed by WLC physics into its latent representations. The latent representations from both autoencoders are then jointly optimized for clustering assignments through a gated fusion mechanism and spectral K-means loss. Further details on all methods are provided in the Section *Methods*.

**Figure 2:**
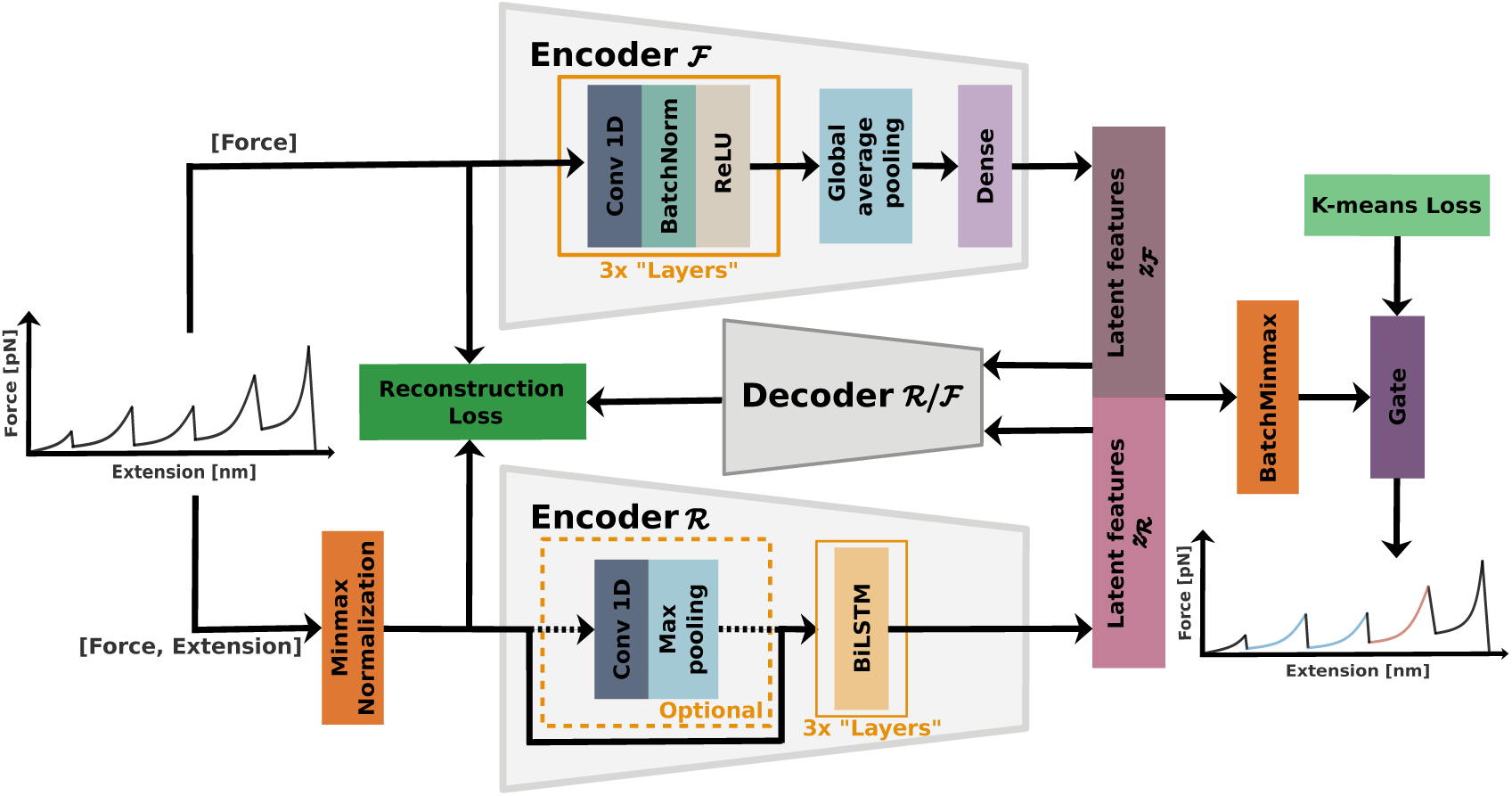
**Model architecture of LatentUnfold**, a novel physics-aware deep clustering framework.

### Validation on Simulated Data

To validate our clustering methods, we employed our Monte Carlo simulation engine, ^31^ based on the Worm-Like Chain (WLC) model ^32^ and the Dudko–Hummer–Szabo model, ^33^ to generate synthetic datasets with known ground truth. We then evaluated performance using clustering accuracy and the Silhouette score. Clustering accuracy ^34^ is an external metric that measures how well predicted cluster labels align with ground truth labels. The Silhouette score ^35^ is an internal metric that assesses cluster compactness and separation without requiring ground truth. Specifically, it quantifies how similar each unfolding segment is to its assigned cluster compared to the other clusters, where higher scores indicate more compact and well-separated clusters. We further validate our clustering results by examining two physical properties that quantify mechanical behavior: ^1^ the unfolding force, defined as the peak force within an unfolding segment, and the contour length increase (Δ*L_c_*), defined as the difference in WLC-fitted contour length (*L_c_*) (Equation 7) between consecutive unfolding segments. Each dataset produced approximately 1,000 simulated force-extension curves at a fixed pulling speed. We tested all clustering methods across 51 simulated datasets (17 parameter configurations *×* 3 pulling speeds: 500, 1000, and 2000 nm/s). A systematic robustness study confirmed that predicted distributions remained closely aligned with ground truth across a wide range of parameter perturbations, and an ablation study demonstrated that both the force and relationship autoencoders are necessary for optimal performance in LatentUnfold. Full simulation results are provided in the *Supporting Information*. We next evaluate our methods on the synthesized protein molecule ddFLN4–Titin I27, and then extend the analysis to natural protein molecules, including bact UtrN-R3, insect UtrN-R3, and DysN-R3.

### Recovering Known Domain Properties in ddFLN4–Titin I27

Unlike simulated data, real experimental data lacks ground-truth labels and may contain traces that do not originate from single molecule events. As a result, we use only the Silhouette score as the evaluation metric for experimental data. To ensure data quality, we apply a two-step filtering process, with further details provided in the Methods: 1) apply a physics-augmented deep learning model for initial filtering to remove traces not originating from single molecule proteins; ^31^ and 2) select unfolding segments based on the fitting quality to the WLC model. Since clustering labels are arbitrary, we assign label 0 to the cluster with the smaller median unfolding force for consistent presentation of results.

AFM experiments on ddFLN4–Titin I27 were conducted across multiple sessions, with each session performed at a fixed pulling speed of either 500, 1000, or 2000 nm/s, yielding approximately 3,000–5,000 trials per session. To determine the appropriate number of clusters, we evaluated the Silhouette score for different cluster counts *K* (Table 1). When *K* = 1, the Silhouette score is negative, indicating a lack of meaningful clustering structure. The score peaks at 0.65 for *K* = 2, and subsequently decreases as *K* increases; suggesting the presence of two well-separated clusters in the dataset.

**Table 1:** Silhouette scores (mean and standard deviation across five runs and three pulling speeds) for the LatentUnfold(LSTM) model with different cluster numbers on four protein molecules. Additional results and detailed trends are shown in Figure S3 in Supporting Information.

| Molecule | K=1 | K=2 | K=4 |
| --- | --- | --- | --- |
| ddFLN4–Titin I27 | -0.09 (0.23) | **0.65 (0.04)** | 0.51 (0.04) |
| Insect UtrNR3 | -0.08 (0.32) | **0.67 (0.11)** | 0.53 (0.04) |
| Bact UtrNR3 | -0.08 (0.21) | **0.62 (0.07)** | 0.52 (0.03) |
| DysNR3 | -0.12 (0.18) | **0.61 (0.06)** | 0.52 (0.03) |

We applied the top five clustering algorithms—LatentUnfold(CNN-LSTM), LatentUnfold(LSTM), K-means(DTW), K-medoids(DTW), and MLP-AE, as identified from simulation evaluation—independently to the experimental dataset. Because the primary goal of this work is to obtain reliable biological conclusions about domain-level mechanical heterogeneity rather than to benchmark a single algorithm, we adopted a consensus voting strategy among the five methods: only unfolding segments assigned to the same cluster by a majority of methods are included in the reported statistics. This approach increases confidence in the domain assignments by retaining only those results that are robust across methodologically diverse algorithms. We note that each method individually produces results consistent with the consensus (see Figure 3 in Supporting Information), further supporting the reliability of the findings.

**Figure 3:**
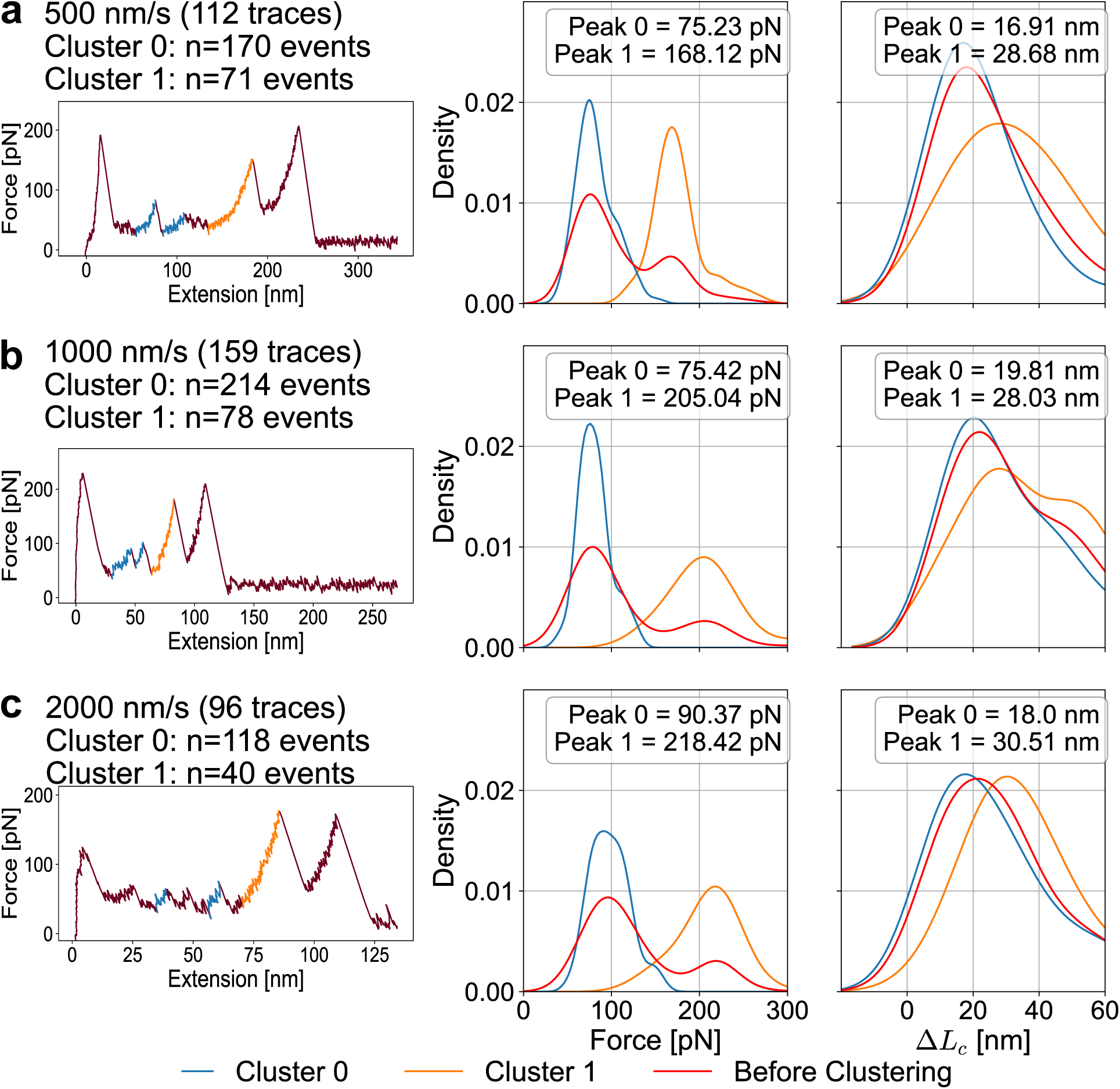
Clustering results for a synthesized protein with well-characterized domains. Clustering statistics for ddFLN4 (cluster 0) and Titin I27 (cluster 1) at pulling speeds of (a) 500 nm/s, (b) 1000 nm/s, and (c) 2000 nm/s. The first column shows representative force-extension curves with color-coded segments by predicted cluster labels. The second column displays the unfolding force distributions, and the third column presents the distributions of contour length increase (Δ*L_c_*). Cluster 0 is shown in blue, Cluster 1 in orange, and the combined distribution is shown in red.

After filtering, Each pulling speed condition (500, 1000, and 2000 nm/s) includes more than 95 single-molecule force-extension curves, with each predicted cluster containing at least 40 unfolding events. According to previous literature, ddFLN4 unfolds via a two-step pathway with unfolding forces of around 60-80 pN at pulling speeds of 1000 nm/s, and the contour length increase of *∼*16 nm per step. ^36^ Titin I27 unfolds in a single step, with unfolding forces of 200–220 pN and a contour length increase of *∼*28 nm. ^3,37^ In our clustering results (Figure 3), cluster 0 is assigned to the group with the lower median unfolding force, corresponding to ddFLN4, while cluster 1 represents Titin I27. At 1000 nm/s, Cluster 0 (ddFLN4) exhibits a peak unfolding force of 75 pN, whereas Cluster 1 (Titin I27) peaks at 205 pN. The estimated contour length increases for ddFLN4 and Titin I27 are approximately 20 nm and 28 nm, respectively. These align well with known reported values. As the pulling speed increases from 500 nm/s to 2000 nm/s, the unfolding forces for both clusters increase, while the contour length increases remain largely unchanged. Along with the representative force-extension curves shown in Figure 3b, these results demonstrate that our method successfully identifies heterogeneous domains and accurately attributes them to ddFLN4 and Titin I27.

### Uncovering Mechanically Distinct Domain Properties in Utrophin and Dystrophin

Consistent with the trend observed for ddFLN4–Titin I27, all dystrophin and utrophin constructs (insect UtrNR3, bacterial UtrNR3, and DysNR3) exhibit peak Silhouette scores of at least 0.61 at *K* = 2 (Table 1), indicating the presence of two mechanically distinct domain populations. This is consistent with the known structural distinction between the NT domain and the SLRs. Each SLR typically adopts a triple *α*-helical bundle conformation with an approximate folded length of *∼*6 nm, ^38,39^ while the folded length of the NT domain is estimated to be *∼*12 nm. ^40^ The theoretical contour length increase (Δ*L_c_*) upon unfolding of a domain can be calculated using the equation ^1,41,42^

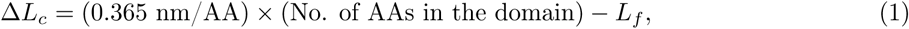

where 0.365 nm/AA is the approximate contour length per amino acid in a fully extended polypeptide chain, and *L_f_* is the folded length of the domain. Thus, the expected difference in contour length increase between the NT domain and a single SLR is approximately 35 nm. This estimate arises from the NT domain having roughly 120 additional amino acids (0.365 *×* 120 = 43.8 nm) compared to an SLR, minus an estimated 6 nm increase in folded length, yielding a net difference of *∼*35 nm.

The analysis pipeline follows the same procedure as described for ddFLN4–Titin I27. Clustering statistics at a pulling speed of 1000 nm/s are presented in Figure 4 for DysNR3, bacterial UtrNR3, and insect UtrNR3, from top to bottom, alongside representative force-extension curves. Results for 500 and 2000 nm/s are provided in Figure S5 in Supporting Information. Across all pulling speeds and constructs, Cluster 1 consistently exhibits higher contour length increases Δ*L_c_* and higher unfolding forces compared to Cluster 0. This observation supports our assumption that Cluster 1 corresponds to the NT domain, while Cluster 0 represents the SLRs. The difference in Δ*L_c_* between the two clusters ranges from approximately 18 to 42 nm across all speeds and constructs, bracketing the theoretical estimate of *∼*35 nm. The variability in observed Δ*L_c_* differences likely reflects partial unfolding events, variability in WLC fitting, and differences in unfolding pathways across experimental conditions. For DysNR3, the most probable unfolding force differences between clusters are 68 pN at 500 nm/s, 91 pN at 1000 nm/s, and 73 pN at 2000 nm/s. Furthermore, unfolding forces increase with higher pulling speeds across all three constructs, consistent with biophysical behavior.

**Figure 4:**
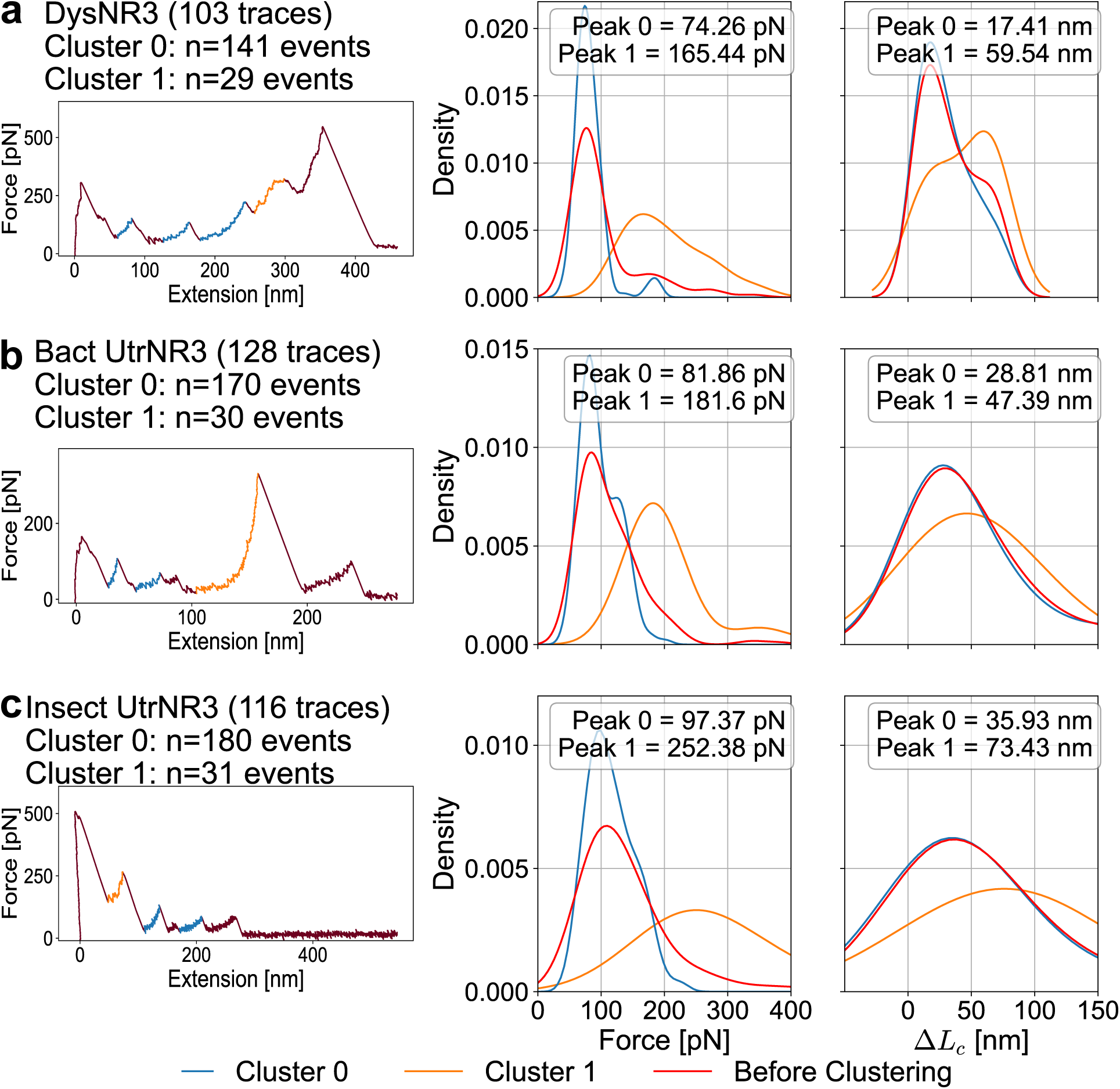
Results of natural protein molecules. Clustering statistics at 1000 nm/s for (a) DysNR3, (b) Bact UtrNR3, and (c) Insect UtrNR3.

Finally, we examined the unfolding order of three natural protein molecules—insect UtrN-R3, bacterial UtrN-R3, and DysN-R3. As summarized in Table 2, three dominant unfolding patterns emerge: (1) SLR domains unfold prior to the NT domain (SLRs *→* NT), (2) the NT domain unfolds between SLR domains (SLRs *→* NT *→* SLRs), and (3) the NT domain unfolds before the SLR domains (NT *→* SLRs). Here, SLRs denotes one or more SLR domains unfolding. Across all three molecules, the most probable pathway (68%) is that the NT domain unfolds after all SLR domains have unfolded. In contrast, there is a moderate probability (25–29%) that the NT domain unfolds before the SLRs. The intermediate scenario, in which the NT domain unfolds between SLR segments, is comparatively rare (4–7%).

**Table 2:**
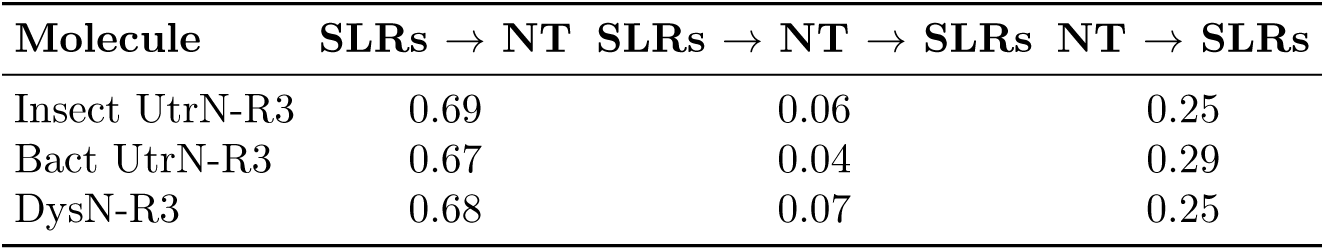
Probabilities of unfolding order for natural protein molecules, insect UtrN-R3, bacterial UtrN-R3, and DysN-R3. SLRs denotes one or more SLR domains unfolding.

These results suggest a clear mechanical hierarchy among domains. SLR regions, composed of repetitive *α*-helical motifs, are generally more mechanically labile, making them prone to unfolding at lower forces. In contrast, the NT domain is more mechanically stable, allowing it to resist unfolding until higher forces are reached. This explains why the NT domain most frequently unfolds last. The low probability of interleaved unfolding further indicates that force propagation favors a sequential, stability-driven pathway rather than frequent alternation between domains. Notably, these observations are enabled by our clustering-based analysis framework and provide, to our knowledge, the first quantitative characterization of unfolding order probabilities in both dystrophin and utrophin.

## Conclusions

In this study, we provide the first automated approach for identifying heterogeneous mechanical domains from SMFS data, leveraging both existing clustering models and a novel physics-aware deep clustering architecture, LatentUnfold. We validated our framework on a synthetic protein composed of two well-characterized domains (ddFLN4–Titin I27), showing that the recovered mechanical properties closely match previously reported values. We then applied our approach to natural protein constructs of dystrophin and utrophin, successfully uncovering two mechanically distinct domain populations with different unfolding forces and contour length increases.

The identification of heterogeneous domains in the dystrophin and utrophin constructs carries direct biological implications. Our clustering framework resolved unfolding segments in all three natural constructs— insect UtrN-R3, bacterial UtrN-R3, and DysN-R3—into two distinct populations: a higher-force, higher-Δ*L_c_* cluster attributed to the NT domain and a lower-force cluster attributed to the SLRs. Across all molecules, the NT domain predominantly unfolds after all SLRs, with interleaved unfolding being rare (Table 2), establishing a clear mechanical hierarchy: the SLRs yield first, while the more stable NT domain persists until higher forces are reached. Force propagation thus follows a sequential, stability-driven pathway rather than alternating between domain types. Such domain-level resolution was previously inaccessible when mechanical properties could only be assessed as averages across all domains. This distinction is particularly relevant given that dystrophin is hypothesized to function as a molecular shock absorber, in which the spectrin-like repeats unfold under mechanical stress to dissipate energy and protect the sarcolemma. ^10^ The differential force response of the NT domain and SLRs revealed here provides a more complete picture of how dystrophin absorbs mechanical energy during muscle contraction and stretching.

These findings also have implications for Duchenne Muscular Dystrophy therapeutics. Utrophin is under active investigation as a potential replacement for dystrophin in DMD, yet recent work has shown that dystrophin and utrophin exhibit fundamentally different mechanical behaviors at the whole-molecule level. ^5^ Our domain-level analysis adds a new dimension to this comparison: the mechanical heterogeneity within each protein can now be characterized, enabling more detailed assessments of whether individual domains of utrophin can functionally substitute for their dystrophin counterparts.

The present study focused on proteins with two mechanically distinct domain types (*K* = 2), which is supported by the Silhouette score analysis across all four protein systems. Many natural proteins, however, contain a larger number of heterogeneous domain types. Extending this framework to *K >* 2 is a natural next step, though it will require careful consideration of how to determine the optimal number of clusters and how to validate assignments for more complex domain architectures.

## Methods

### Protein preparation

The synthetic protein ddFLN4-Titin I27 was expressed and purified from the plasmid described in.^43^ DysN-R3, insect UtrN-R3, and bact UtrN-R3 were cloned, expressed, and purified as previously described. ^6,29^

### Atomic force microscopy experiments

All SMFS experiments were performed using an MFP-3D atomic force microscope (Asylum Research, Oxford Instruments, Santa Barbara, CA) equipped with a laser-photodiode deflection sensor and a piezoelectric nano-positioner. ^16^ Bruker MLCT-BIO silicon nitride cantilevers with gold-coated backsides (nominal spring constant 10 pN/nm, typical tip radius 20 nm) were used throughout. The spring constant of each cantilever was calibrated by thermal noise analysis prior to each experiment.^44^

Purified protein in phosphate-buffered saline (PBS) was deposited at 3–30 nMol onto freshly cleaved, ionized mica substrates and incubated for 15 minutes to promote surface adhesion. The substrate was then washed with 100 *µl* PBS to remove unbound protein, and a fresh 100 *µl* PBS droplet was added for the measurements. Protein concentrations were chosen to maintain a 5–10% probability of successful single-molecule pulls, consistent with established protocols for minimizing multi-molecule events.^5,45^

Force spectroscopy was conducted at room temperature (21–23°C) using repeated approach-retraction cycles. In each cycle, the cantilever tip was pressed against the substrate for 2 seconds at an indentation force of 600 pN, then retracted at constant speeds ranging from 200 to 5000 nm/s. The force *F* on the protein was determined from the cantilever deflection as

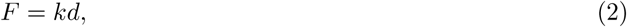

where *k* is the spring constant and *d* is the deflection of the cantilever.

### Experimental data analysis

Because the size of a typical force probe is orders of magnitude larger than that of a single biomolecule, it may come into contact with one or more molecules during an experiment. To reduce the likelihood of multiple molecule interactions, the concentration of biomolecules is typically lowered. ^6,45^ However, this introduces a trade-off: while lower concentrations reduce multi-molecule events, they also increase the frequency of measurements with no molecular interaction. To obtain accurate and interpretable data from single-molecule force spectroscopy (SMFS), it is essential to exclude traces arising from both cases—those containing only background noise due to the absence of a molecule, and those involving multiple molecules, which introduce confounding interpretation.

To ensure data quality in experimental data, we employed a two-step filtering process. First, we apply a physics-augmented deep learning model ^31^ to pre-filter experimental data and identify those most likely originating from single-molecule measurements. Although deep learning models are not perfectly accurate, this initial screening substantially reduces the number of traces requiring manual inspection. Next, we filter unfolding segments based on their fitting quality to the worm-like chain (WLC) model. Unfolding segments are first identified by detecting characteristic peaks in the force signal that exceed a threshold of 3*σ*, where *σ* is the standard deviation of noise. The first and last significant events typically arise from adhesive forces and detachment events, respectively, and are excluded from further analysis. The remaining significant peaks are interpreted as domain unfolding events of interest. These segments are then fitted to the WLC model, as defined in Equation 7, and only those with high fitting quality are retained for downstream analysis.

### LatentUnfold

The force encoder *f_encF_* maps *F*^(^*^i^*^)^ into a latent representation 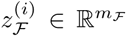 with *m_F_ ∈* N^+^ and the decoder *f_decF_* reconstructs the data from the latent space. After both encoding and decoding process, the reconstructed force sequence *F*^^(^*^i^*^)^ *∈* R*^T^* is computed as *F*^^(^*^i^*^)^ = *f_dec_*(*f_encF_*(*F*^(^*^i^*^)^)).

We use the mean square error (MSE) as the reconstruction loss to train the autoencoder with *Y* being the input and *Y*^^^ being the reconstruction, defined as:

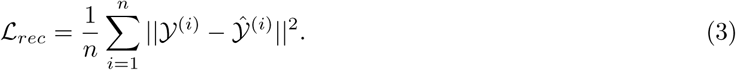

The force encoder architecture consists of three convolutional blocks, followed by a global average pooling layer and a fully connected layer to produce the latent representation. Each convolutional block is composed of a 1-Dimensional convolutional layer, batch normalization layer ^46^ and a Rectified Linear Unit (ReLU) ^47^ activation layer. The decoder is constructed as the mirror of the encoder architecture with three convolutional blocks.

The relationship encoder *f_encR_* extracts a latent representation 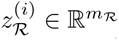 with *m_R_ ∈* N^+^, which is then passed through a relationship decoder *f_decR_* to reconstruct both sequences (*F*^^(^*^i^*^)^*, X*^^(^*^i^*^)^). This autoencoder is trained using the reconstruction loss defined in Equation 3. We tested two encoder architectures, the first one, referred to as LatentUnfold(LSTM), uses three stacked bidirectional long short-term memory (BiLSTM) layers ^48^ with the decoder is constructed as the mirror of the encoder architecture. The second, referred to as LatentUnfold(CNN-LSTM) inspired by deep temporal clustering, ^49^ introduces a convolutional layer followed by a max-pooling layer prior to the BiLSTM stack to better capture cross-channel relationships, corresponding to the optional block in Figure 2.

After obtaining the two latent representations *z_F_* and *z_R_*, we apply feature-wise min-max normalization to remove scale differences between them. To adaptively control the influence of each representation on the clustering task, we introduce learnable parameters *α_F_* and *α_R_*, computed as *α_F_, α_R_* = *softmax*(*W ·* [*z_F_ z_R_*] + *b*), with *W ∈* R*^n×^*^(^*^mF^* ^+^*^mR^*^)^ and *b ∈* R*^n^* are weights and bias of gate block.

These weights modulate the relative importance of each latent space. The final combined representation is *H* = [*α_F_ · z_F_* ; *α_R_ · z_R_*] *∈* R^(^*^mF^* ^+^*^mR^*^)^*^×n^*. We optimize a spectral-relaxed K-means loss ^27,50^ defined as:

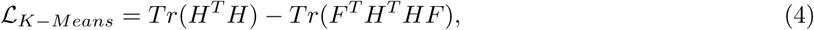

where *Tr*(*·*) denotes the matrix trace and *F ∈* R*^n×K^* is the cluster indicator matrix, where *K* is the number of clusters.

### State-of-the-art clustering methods

K-Means is a widely used clustering algorithm that assigns data points to the nearest centroid to minimize intra-cluster variance:

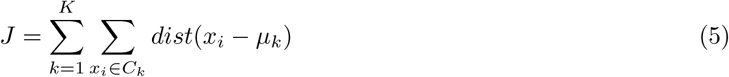

where *K* is the number of clusters, *x_i_* is a data point, *µ_k_* is the centroid of cluster *C_k_* and *dist*(*·*) denotes the distance. K-Medoids selects actual data points (medoids) as cluster centers, making it more robust to noise and outliers. Common options for distance are Euclidean norm and DTW. However, applying it directly to time series data using Euclidean norm is inadequate due to temporal misalignments. DTW addresses this by computing the minimum cumulative distance between aligned points of two sequences *X* = [*x*_1_*, x*_2_*, …x_n_*] and *Y* = [*y*_1_*, y*_2_*, …y_m_*], allowing for elastic shifts in the time axis.

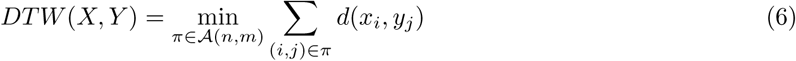

where *π* is a warping path from start to end, *A*(*n, m*) is the set of all valid warping paths and *d*(*x_i_, y_j_*) is the distance function (usually Euclidean distance).

Autoencoders (AEs) are unsupervised neural networks designed to learn low-dimensional latent representations by reconstructing the input. ^25^ For instance, MLP-AE consists of fully connected layers, and ResNet-AE uses multiple convolutional layers with residual connections, followed by a global average pooling and a dense embedding layer. These autoencoders are trained solely with reconstruction loss, and clustering is assigned with the learned latent representations using K-Means.

DEC ^26^ is a deep clustering framework where a neural network is trained to jointly learn latent representations and cluster assignments. Deep temporal clustering representation (DTCR)^27^ is a deep clustering method tailored for time series data. It uses stacked bidirectional gated recurrent units (GRUs) to encode temporal patterns and integrates three objectives: reconstruction loss, a binary classification loss to distinguish real from time-shuffled inputs, and a K-means loss to learn cluster assignments.

## Dataset

### Simulation data

A Monte Carlo simulation based method, adopted from, ^31^ was used to generate simulated force-extension data. The system consists of a single protein composed of two distinct domain types, indexed by *i* = 0, 1. The cantilever base is pulled at constant velocity *v*, and at each time step Δ*t* the protein extension *X* is calculated via a force-balance equation using the Worm-Like Chain (WLC) model: ^32^

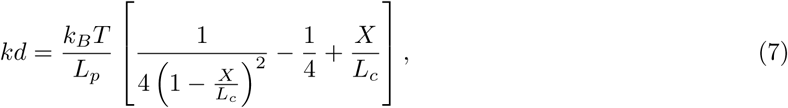

where *k_B_*is the Boltzmann constant, *T* is temperature, and *L_c_*and *L_p_* are the contour length and the persistence length of the protein, respectively.

At each time step, the probability that a domain of type *i* = 0, 1 unfolds is given by

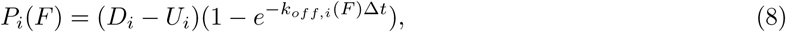

where *D_i_*is total number of domains and *U_i_* is the number of unfolded domains of type *i*, and *k_off,i_*(*F*) is the transition rate that can be determined with the Dudko-Hummer-Szabo model: ^33^

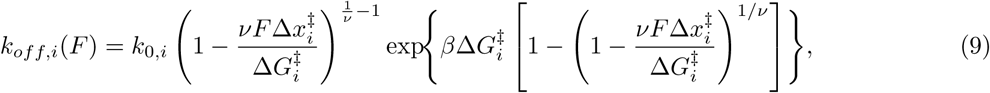

where *k*_0*,i*_ is the intrinsic transition rate, 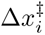 is the distance to energy barrier, 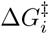 is the energy barrier height for type *i* domain, and 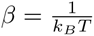, and *ν* = 1*/*2 or 2*/*3, representing the cusp-like or linear-cubic energy landscape.

To determine whether unfolding occurs, a random number between 0 and 1 is generated and compared to *P_i_*(*F*). If the random number is smaller than *P_i_*(*F*), the type *i* domain unfolds. If both *P*_0_(*F*) and *P*_1_(*F*), exceed the random number, the domain with the higher probability unfolds, while the other remains folded. Upon unfolding, the contour length of the entire protein is updated by adding an increment Δ*L_c,i_* corresponding to the unfolded domain type. The two domain types differ in both their kinetic parameters, *k*_0_, Δ*x^‡^*, and Δ*G^‡^*, and their elastic property such as the contour length increment Δ*L_c_*.

### Experimental data

Experimental SMFS data were obtained from AFM experiments on four protein molecules, as described in the Introduction: the synthetic two-domain protein ddFLN4–Titin I27, and three natural protein constructs DysN-R3, insect UtrN-R3, and bact UtrN-R3. The two utrophin constructs differ in expression system (insect vs. bacterial cells), which has been shown to influence mechanical properties. ^6,29^

### Training details and evaluation metrics

All clustering models, including LatentUnfold, are trained independently on each protein system in a fully unsupervised manner without any domain labels. This per-protein training strategy means that no single trained model is transferred across protein systems; rather, the framework is applied de novo to each dataset, and its generalizability is demonstrated by consistent performance across four structurally diverse protein molecules.

In our model, the force autoencoder employs convolutional layers with a fixed filter size of 64. The filter lengths for the first, second, and third convolutional layers are set to 8, 5, and 3, respectively. The default latent space dimension for the force encoder is *m_F_* = 2. For the relationship autoencoder, three stacked BiLSTM layers uses size of 64,8,2, respectively, resulting in a latent space dimension of *m_R_* = 4. In the CNN-LSTM variant, a convolutional layer with 100 filters of size 10 is applied before the BiLSTM layers, followed by a max-pooling layer of size 10. All models are trained with a batch size of 16 for 50 epochs. All models are trained with Adam ^51^ with the learning rate 0.001, *β*_1_ = 0.9, *β*_2_ = 0.999 and *ɛ* = 1*e −* 8. Experiments are conducted on a high-performance computing setup featuring NVIDIA and Apple M1 Pro GPUs, ensuring computational efficiency. All models, including LatentUnfold and the baseline methods, require less than one hour for both training and inference on each dataset, making the framework practical for routine SMFS analysis. Each dataset is partitioned into training and test subsets using a 20-80 split.

Clustering accuracy ^34^ is an external metric that measures how well the predicted cluster labels *y_pred_* align with the ground truth labels *y_true_*. Since cluster labels are arbitrary, a permutation function *π*(*·*) is applied to find the best label alignment. The accuracy is defined as

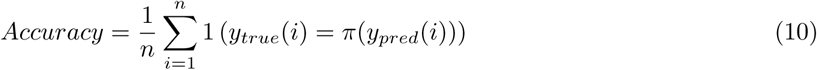

where *n* is the number of samples and 1(*·*)is the indicator function. As an internal metric, the Silhouette score ^35^ assesses the compactness and separation of the clusters without using ground truth. For each sample *i*, *a*(*i*) is the average distance to other points in the same cluster, and *b*(*i*) is the minimum average distance to points in any other cluster. The Silhouette score is defined as

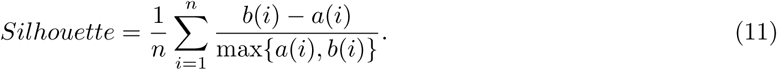

Higher values indicate better-defined c lusters, w ith 1 b eing i deal s eparation a nd v alues n ear 0 indicating overlapping clusters.

## Acknowledgements

This project was supported by funding from NIH (5R01AR042423).

## Supporting information

### Results on simulation data

Two widely used physical properties are the unfolding force and the contour length increase. ^1^ The unfolding force, the peak force within an unfolding segment, can be described by the Dudko-Hummer-Szabo (DHS) model, ^33^ whose parameters include the intrinsic transition rate *k*_0_, the distance to the energy barrier Δ*x^‡^*, and the energy barrier height Δ*G^‡^*. The contour length *L_c_*, the maximum physically possible extension, is obtained by fitting the WLC model (Equation)); the contour-length increase Δ*L_c_* is the difference in *L_c_* between consecutive unfolding segments.

The simulation data generation process was described in the *Methods* section, where each dataset was created using a distinct pair of parameters. Each dataset produced approximately 1,000 simulated force-extension curves at a fixed pulling speed. We simulated datasets at three pulling speeds: 500, 1000, and 2000 nm/s. Since the ground truth labels are known for the simulated data, we evaluated clustering performance using both accuracy and Silhouette score. Each model configuration is trained and evaluated five times independently to account for stochasticity and ensure statistical robustness.

We first evaluate our approach on simulated datasets generated from two well-separated parameter sets, referred to as ‘P0’ and ‘P1’ (Table 3). As shown in Figure 6a, the predicted unfolding force distributions closely match the ground truth at 1000 nm/s. This agreement persists across other pulling speeds (500 and 2000 nm/s), and similar consistency is observed for contour length increments Δ*L_c_* (Figure 6b). Predicted unfolding forces not only align well with the ground truth but also exhibit a consistent shift toward higher values with increasing pulling speed, while Δ*L_c_* shows minimal dependence on pulling speed—both trends consistent with known biophysical behavior. Example force extension curves (Figure 6c) further illustrate the identification of distinct unfolding segments. Finally, LatentUnfold(LSTM) and LatentUnfold(CNN-LSTM) achieved an average clustering accuracy of 0.93, with Silhouette scores of 0.61 and 0.75, respectively. These physics-aware architectures outperformed existing clustering methods; for instance, LatentUnfold(CNN-LSTM) sur-passed K-means DTW by 0.03 in accuracy and 0.17 in Silhouette score (Table 4).

**Figure 5:**
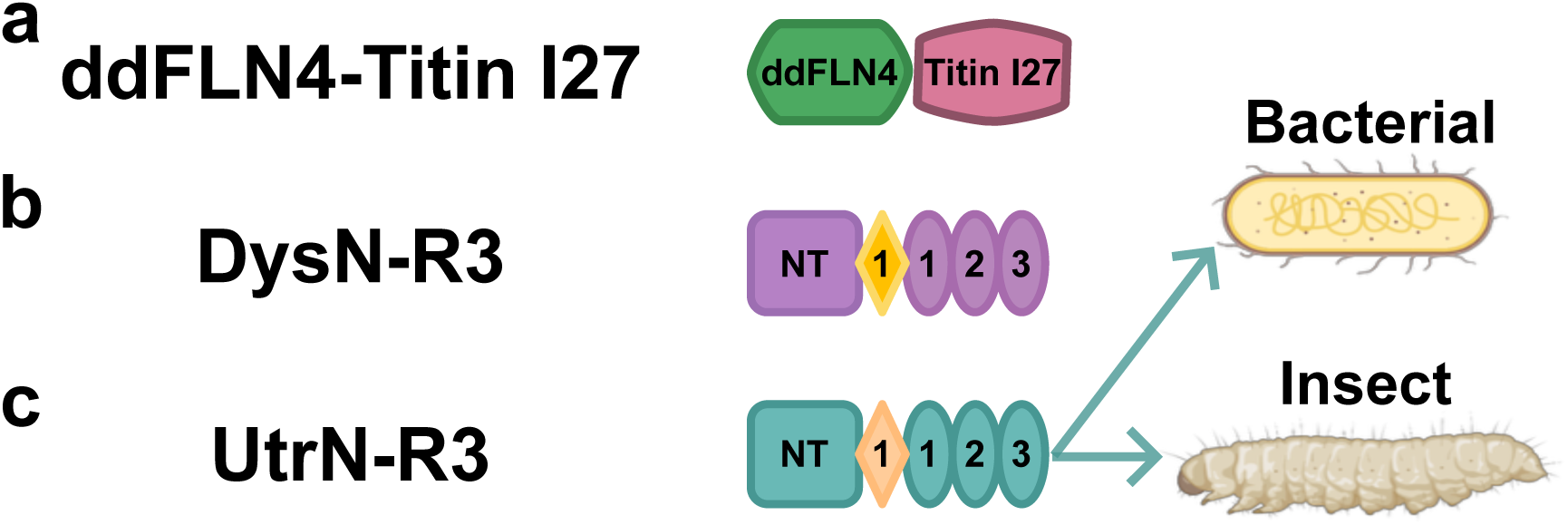
Diagrams of protein molecules. (a) ddFLN4–Titin I27, consisting of one ddFLN4 domain and one Titin I27 domain. (b) DysN-R3, a fragment of dystrophin spanning the N-terminus (NT) through spectrin-like repeat 3 (SLR3). (c) UtrN-R3, a corresponding utrophin fragment, further divided into bacterial UtrN-R3 and insect UtrN-R3 based on the expression system. Ovals represent spectrin-like repeats (SLRs), and NT indicates the N-terminus domain.

**Figure 6:**
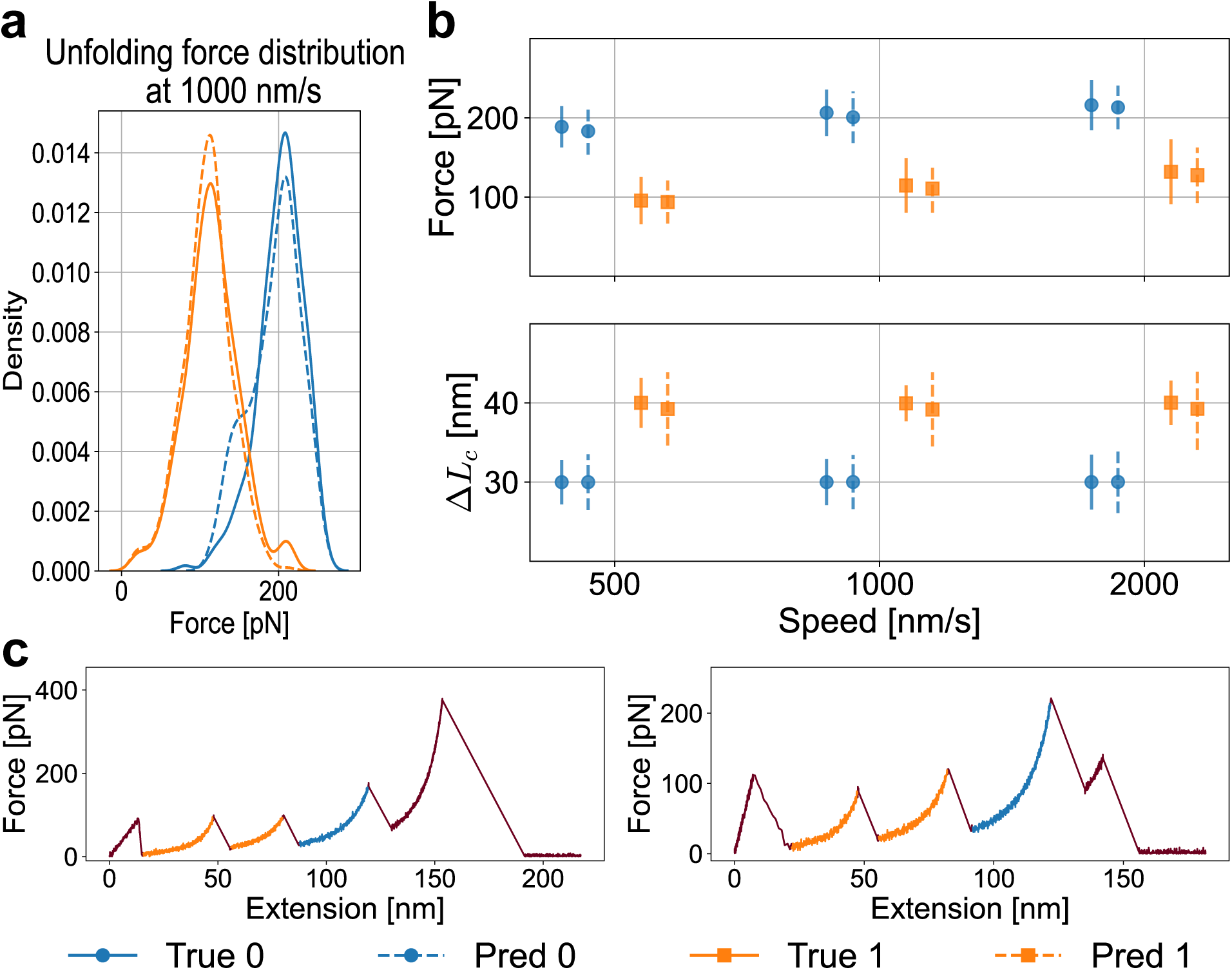
Predicted versus ground truth results on simulated data using LatentUnfold(LSTM). (a) Predicted unfolding force distributions (dashed lines) closely match the ground truth distributions (solid lines) for the dataset at 1000 nm/s. (b) Distributions of unfolding force (top) and contour length increment (bottom), with medians shown as center points and error bars indicating standard deviations. (c) Example force-extension curves with two distinct unfolding segments identified.

**Table 3:** Simulation parameters for parameter sets 0 and 1.

| | $k_0$ | $\Delta x^\ddagger$ [nm] | $\Delta G^\ddagger$ [$k_B T$] | $\Delta L_c$ [nm] |
| --- | --- | --- | --- | --- |
| P0 | 0.0001 | 0.40 | 20.0 | 30.0 |
| P1 | 0.08 | 0.37 | 10.0 | 40.0 |

**Table 4:** Clustering accuracy and Silhouette score (mean with standard deviation in parentheses) for each model at 1000 nm/s. Results at 500 and 2000 nm/s are similar and therefore omitted for brevity. Best values at this speed are highlighted in bold.

| Model | Accuracy | Silhouette |
| --- | --- | --- |
| K-means DTW | 0.90 (0.01) | 0.57 (0.00) |
| K-medoids DTW | 0.91 (0.01) | 0.56 (0.00) |
| MLP-AE | 0.77 (0.12) | 0.51 (0.03) |
| LatentUnfold(CNN-LSTM) | **0.93 (0.02)** | **0.75 (0.02)** |
| LatentUnfold(LSTM) | **0.93 (0.01)** | 0.61 (0.10) |

### Robustness study with systematic parameter variation

Following our initial evaluation on a well-separated parameter set, we conducted a comprehensive robustness study to assess the generalizability of our method across varying simulation conditions. Specifically, we generated 16 additional datasets by systematically varying one of four key parameters at a time, while keeping the remaining parameters fixed at values of ‘P0’, reported in Table 3. The four parameters are: the intrinsic transition rate *k*_0_, the distance to the transition state Δ*x^‡^*, the energy barrier Δ*G^‡^*, and the the contour length increase Δ*L_c_*.

The corresponding results are presented in Figures 7. For example, in one case we increased *k*_0_ from 0.0001 to 0.01—a 99-fold increase—denoted as +99% *k*_0_. Across all tested conditions, the predicted unfolding force and Δ*L_c_* distributions remained closely aligned with the ground truth, demonstrating the robustness of our method. Additionally, the observed effects of each parameter modification match theoretical expectations. Specifically, a smaller *k*_0_, a smaller Δ*x^‡^*, or a larger Δ*G^‡^* each lead to higher unfolding forces without altering Δ*L_c_*. In contrast, changes in Δ*L_c_* primarily affect the contour length increment distribution while having minimal impact on unfolding forces. These results confirm that our clustering framework accurately captures key biophysical properties across a range of controlled parameter perturbations.

**Figure 7:**
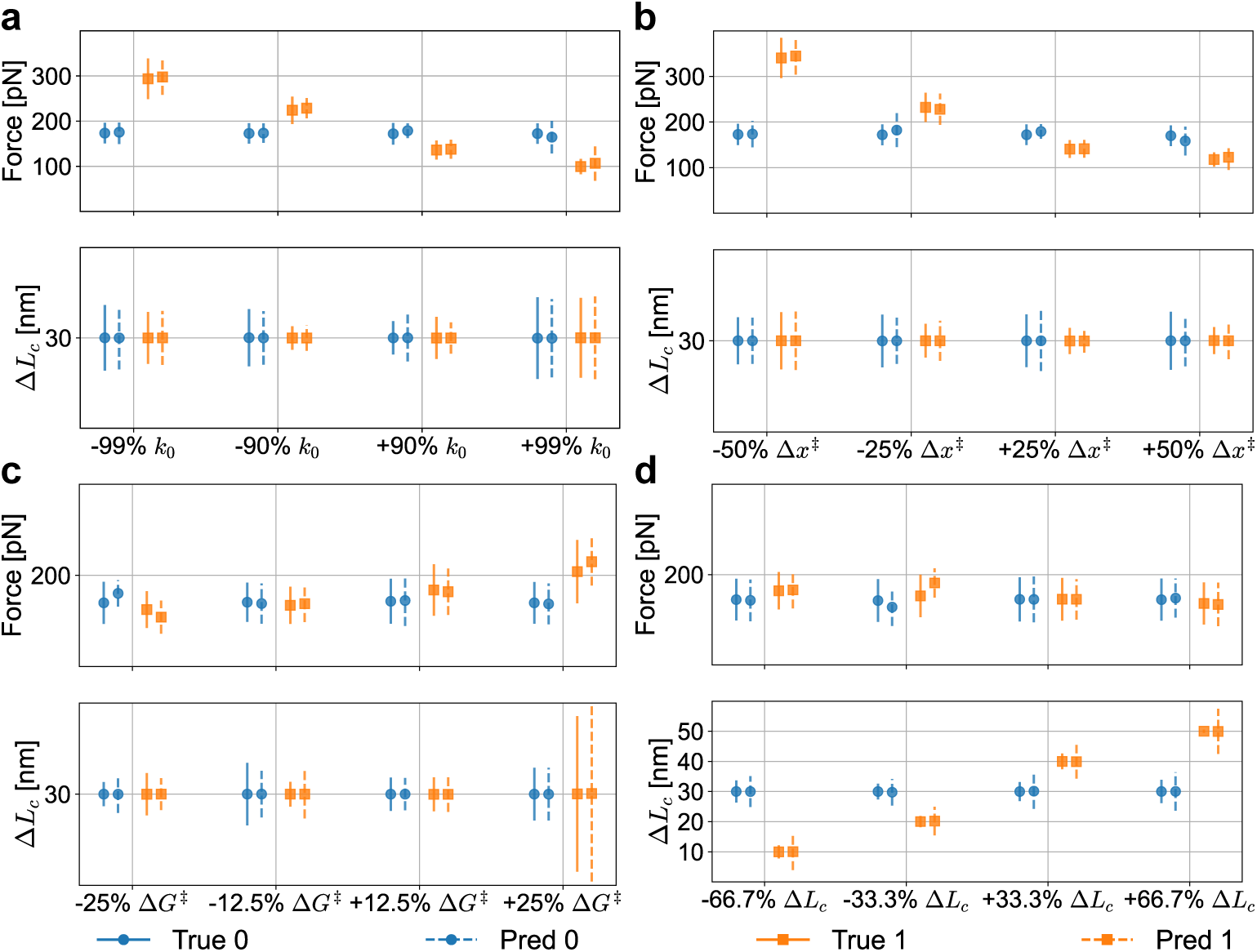
Robustness study under systematic parameter variation. Each parameter is modified by a specified fold relative to values of ‘P0’ (Table 3), and annotated accordingly (e.g., +99% *k*_0_ indicates a 99-fold increase in *k*_0_). Ground truth distributions are shown as solid lines, and predictions as dashed lines.

### LatentUnfold outperforms existing clustering models

We next ranked LatentUnfold with existing clustering models across a total of 51 simulated datasets, corresponding to 17 parameter configurations evaluated at three different pulling speeds. One configuration was generated using the well-separated parameters described above, while the remaining 16 were derived from the robustness analysis detailed in the Supporting Information, where each simulation parameter, *k*_0_, Δ*x^‡^*, Δ*G^‡^*, and Δ*L_c_*, was systematically varied across four values. For each simulation condition, we assessed all models on the corresponding test data and ranked them based on their average performance over five runs using two evaluation metrics: clustering accuracy and Silhouette score. The top-performing model for each parameter pair was assigned a rank of 1, and average rankings were then computed across all parameter combinations. These aggregated results are visualized using critical difference (CD) diagrams. ^52^ As shown in Figure 8a, where clustering accuracy is used as the evaluation metric, our physics-aware models—LatentUnfold(CNN-LSTM) and LatentUnfold(LSTM)—achieved the lowest (i.e., best) average ranks of 2.13 and 3.41, respectively. When Silhouette score is used instead (Figure 8b), LatentUnfold(LSTM) and LatentUnfold(CNN-LSTM) again outperformed, with average ranks of 2.59 and 2.86, respectively. An ablation study (Supporting Information) further confirms that both the force and relationship autoencoders are necessary: variants using only a single autoencoder consistently underperformed the full dual-encoder LatentUnfold architecture.

**Figure 8:**
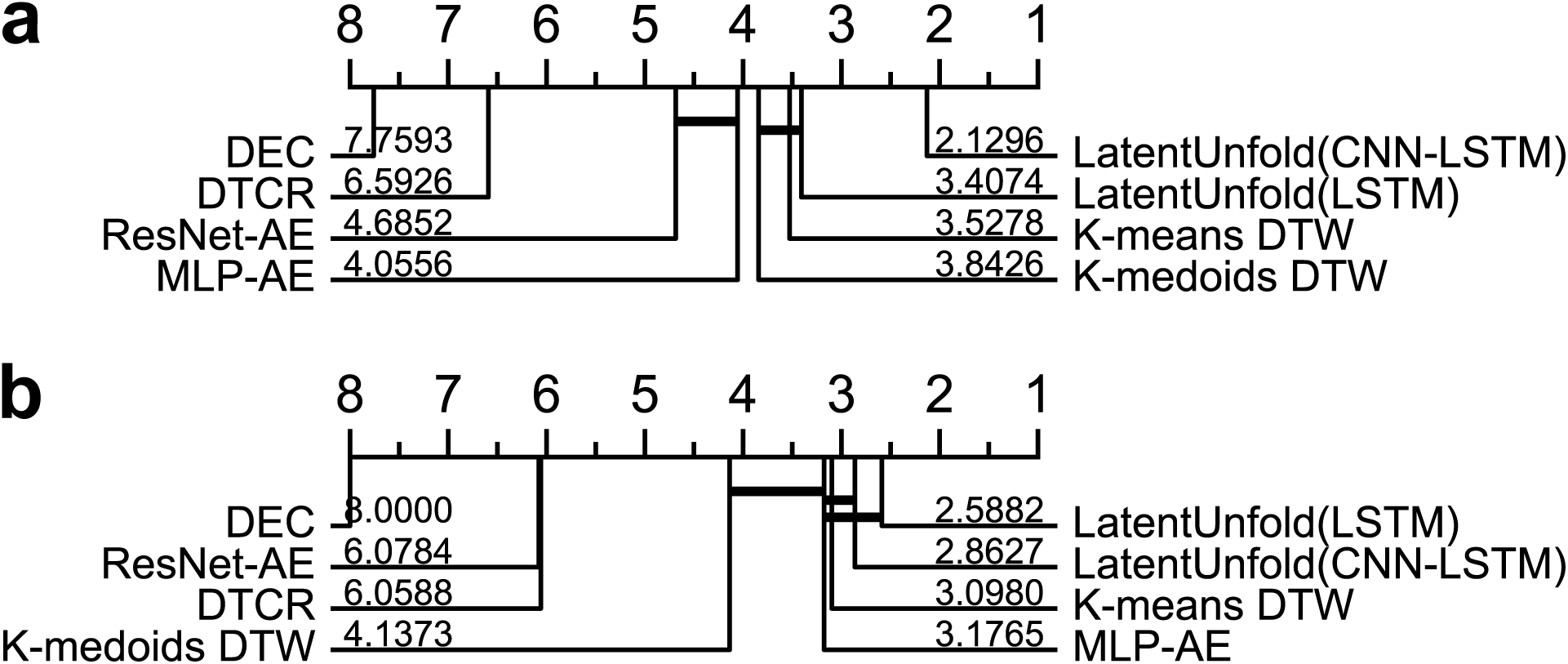
Critical difference (CD) diagrams comparing all models across the 51 simulated datasets using (a) clustering accuracy and (b) Silhouette score. The top-performing model receives a rank of 1. Models connected by a thick horizontal line form a group with no statistically significant difference in performance. Statistical significance is assessed following. ^52,53^

### Ablation study

We conducted an ablation study to evaluate the contribution of different components within our model architecture using all simulated datasets. Our proposed models combine a force autoencoder and a relationship autoencoder, and are further categorized into LatentUnfold(CNN-LSTM) and LatentUnfold(LSTM) based on the architecture of the relationship autoencoder. For comparison, we evaluated three simplified variants that use only a single autoencoder: the force-only autoencoder (FAE), the CNN-LSTM relationship autoencoder (RAE-CNN-LSTM), and LSTM only relationship autoencoder (RAE-LSTM). In these single-autoencoder configurations, clustering is performed by directly applying K-means(DTW) to the learned latent representations. As shown in Figure 9, both LatentUnfold(CNN-LSTM) and LatentUnfold(LSTM) consistently outperform their single-autoencoder counterparts, demonstrating the effectiveness of jointly modeling force and relational information through a dual-autoencoder framework.

**Figure 9:**
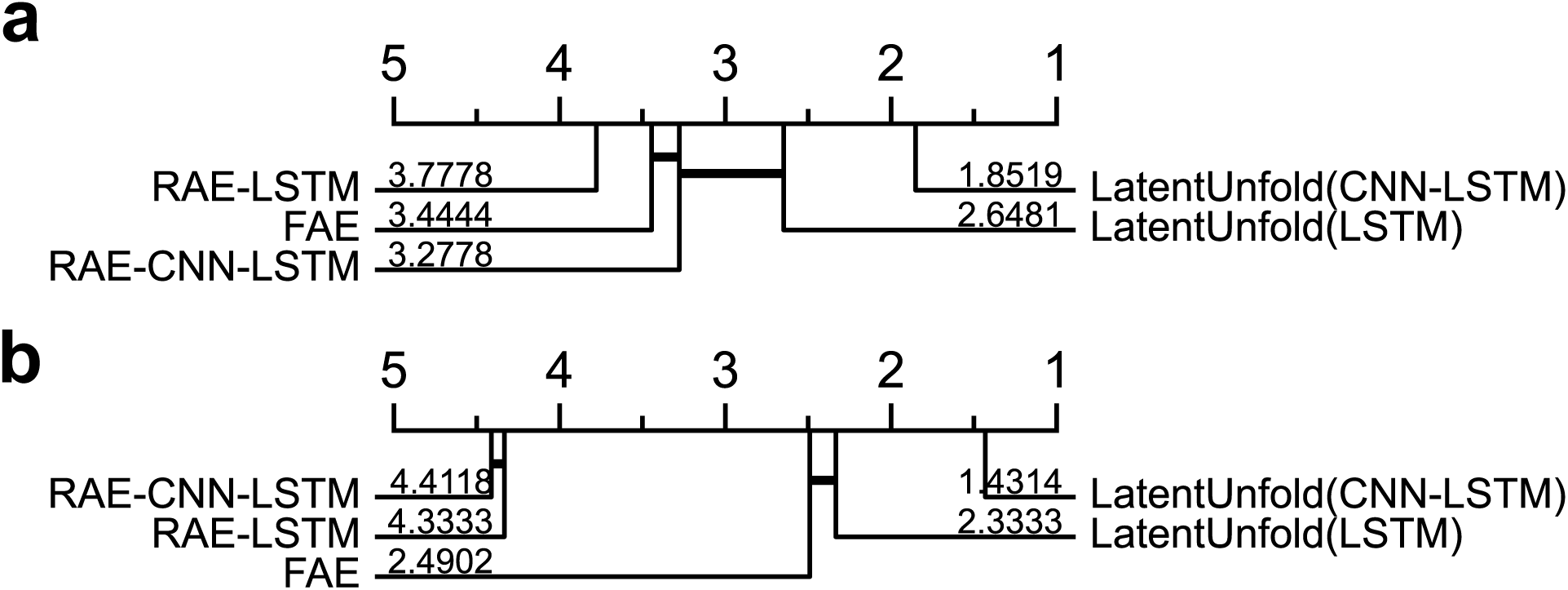
Critical difference (CD) diagram for the ablation study with (a) clustering accuracy and (b) Silhouette score. Both LatentUnfold(CNN-LSTM) and LatentUnfold(LSTM) consistently outperform the single-autoencoder variants: the force-only autoencoder (FAE), the CNN-LSTM relationship autoencoder (RAE-CNN-LSTM), and the LSTM only relationship autoencoder (RAE-LSTM).

## More results on experimental data

### Robustness across top-performing models

We applied five top-performing clustering algorithms—LatentUnfold(CNN-LSTM), LatentUnfold(LSTM), K-means(DTW), K-medoids(DTW), and MLP-AE—independently to the experimental dataset. As shown in Figure 11, the predicted distributions of unfolding force and contour length increase are highly consistent across these models. Furthermore, the level of disagreement in predicted cluster labels, when compared to LatentUnfold(CNN-LSTM), remains relatively low. These findings support the conclusion that our clustering framework yields robust and consistent results across different algorithms, even in the absence of ground truth labels.

**Figure 10:**
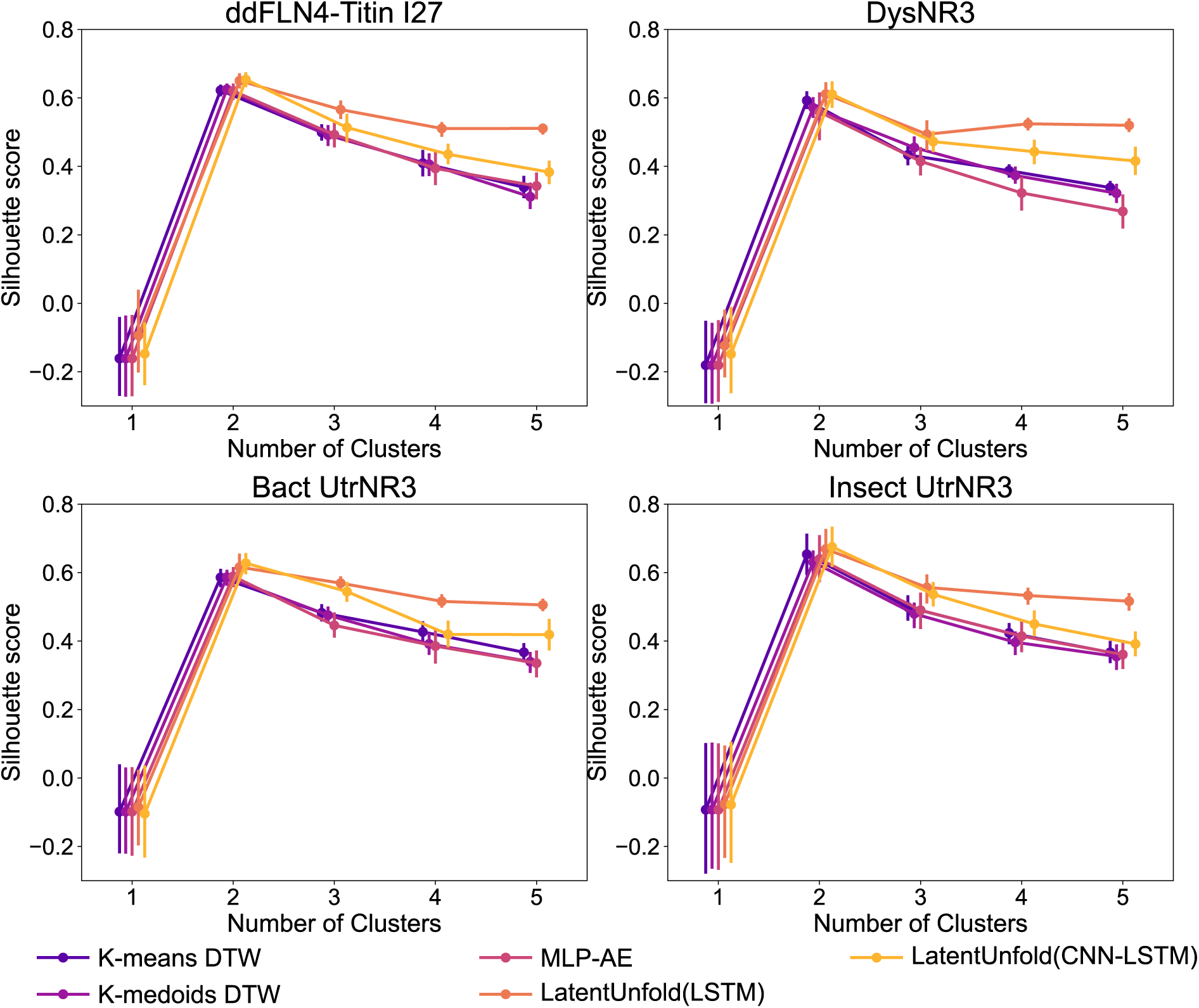
Silhouette score vs number of clusters for all four protein molecules: ddFLN4-Titin I27, bact UtrN-R3, insect UtrN-R3, DysN-R3.

**Figure 11:**
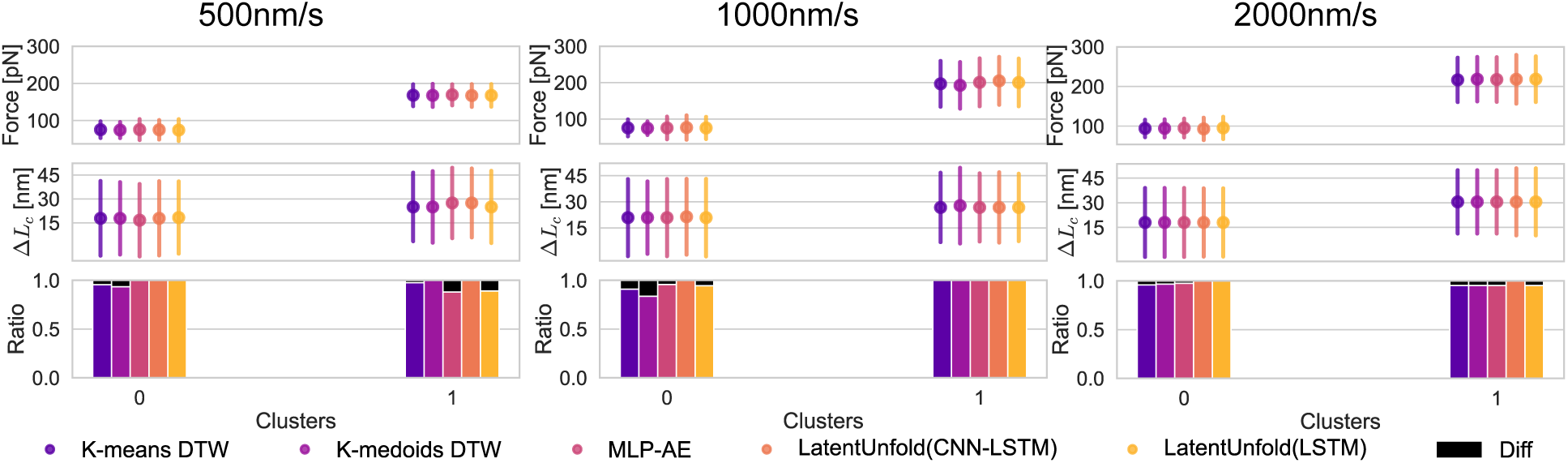
Clustering statistics of ddFLN4–Titin I27 across three pulling speeds: 500, 1000, and 2000 nm/s (from left to right). Dots indicate the most probable values, with error bars representing standard deviations. The first row shows the unfolding force distributions; the second row shows the contour length increase distributions; and the third row presents mismatch ratios, computed by treating the predicted labels from LatentUnfold(CNN-LSTM) as the reference and comparing them against those from the other algorithms. Mismatch ratios are shown in black.

**Figure 12:**
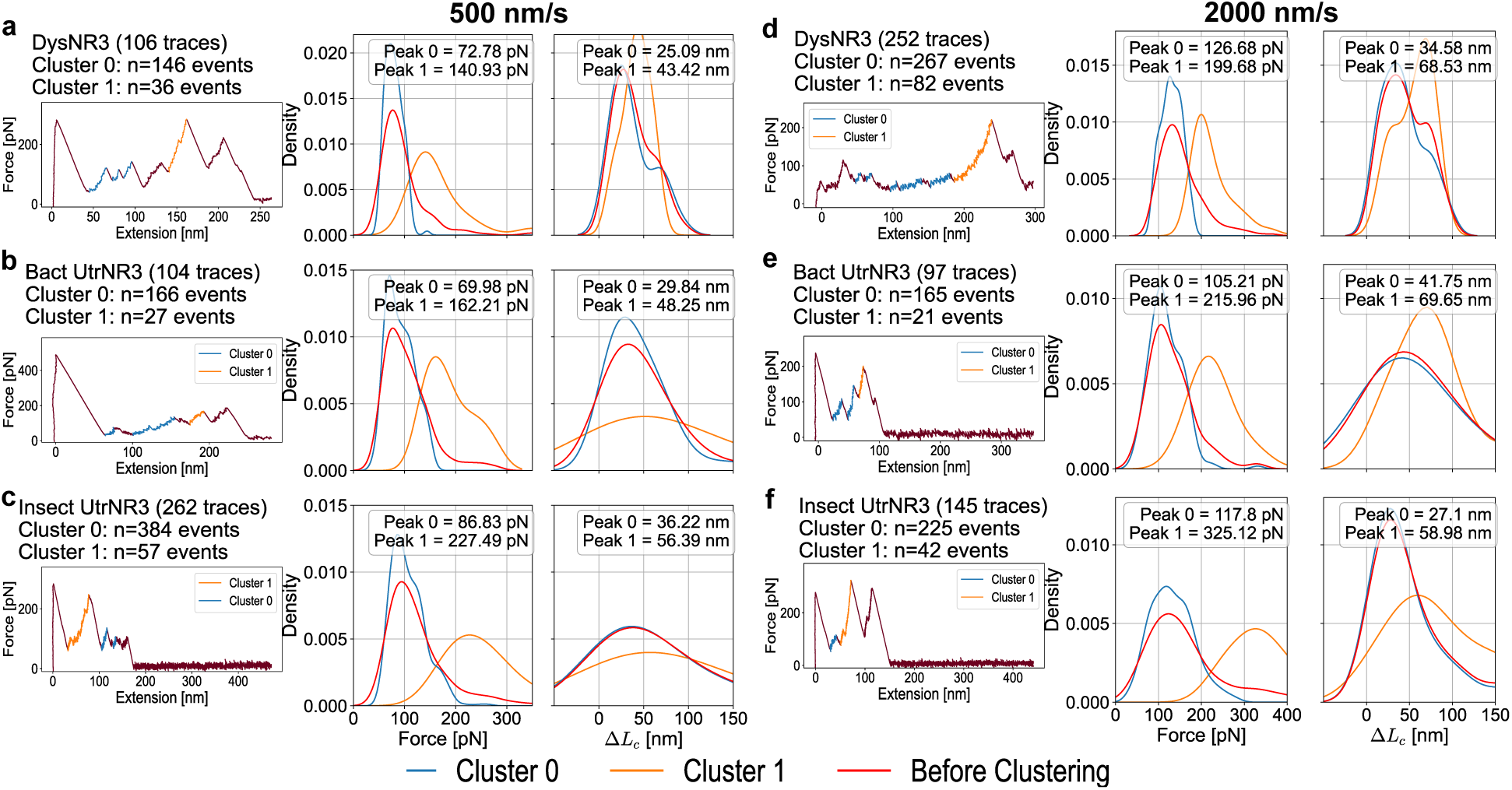
Clustering statistics at 500 nm/s for (a) DysNR3, (b) Bact UtrNR3, and (c) Insect UtrNR3, and at 2000 nm/s for (d) DysNR3, (e) Bact UtrNR3, and (f) Insect UtrNR3. The first column shows representative force-extension curves with color-coded segments by predicted cluster labels. The second column displays the unfolding force distributions, and the third column presents the distributions of contour length increase (Δ*L_c_*). Cluster 0 is shown in blue, Cluster 1 in orange, and the combined distribution is shown in red.

